# Endothelial CEPT1 Regulates Hepatic MTTP-Mediated Lipid Metabolism and Impacts Aortic Atherosclerosis

**DOI:** 10.1101/2025.09.21.677657

**Authors:** Dina Ibrahim, Shahab Hafezi, Tariq Khan, Ibrahim Kuziez, Larisa Belaygorod, Chao Yang, Connor Engel, Chenglong Li, Xiaohua Jin, Maxwell Braasch, Zhengyan Zhang, Yiing Lin, Sangeeta Adak, Clay F. Semenkovich, Mohamed A. Zayed

## Abstract

**Background:** The regulation of hepatic lipid metabolism by vascular endothelial factors remains poorly characterized, despite its relevance to atherosclerosis and steatosis. Microsomal triglyceride transfer protein (MTTP) is essential for hepatic lipid metabolism, but its regulation by endothelial cells has not been previously investigated.

**Objective:** This study examined whether endothelial choline ethanolamine phosphotransferase 1 (CEPT1) modulates hepatic MTTP activity, impacting systemic lipid homeostasis and aortic plaque formation.

**Methods and Results:** Human steatotic liver samples exhibited reduced CEPT1 and MTTP protein levels, correlating with diminished lipid exports. In mice, endothelial-specific *Cept1* knockdown decreased hepatic MTTP expression, reduced serum triglyceride and cholesterol levels, and markedly attenuated aortic atherosclerosis without evidence of fat malabsorption. *In vitro*, endothelial *CEPT1* silencing suppressed MTTP activity in co-cultured hepatocytes via a paracrine mechanism involving peroxisome proliferator-activated receptor α (PPARα) signaling, which was rescued by fenofibrate treatment. Aortic histology confirmed reduced plaque burden and macrophage infiltration in CEPT1-deficient mice.

**Conclusions:** Endothelial CEPT1 critically regulates hepatic MTTP through a paracrine axis, influencing lipid metabolism and atherogenesis. Targeting endothelial CEPT1 may represent a novel therapeutic approach to reduce steatosis and vascular atherosclerosis.

## Introduction

Globally, it is estimated that 39% of adults (>2 billion individuals) currently suffer from hyperlipidemia leading to atherosclerosis and hepatic steatosis ^1^. The prevalence of these co-morbidities is increasing ^2^. While lipid lowering medications can effectively reduce serum lipid levels, individuals with hepatic steatosis are still at higher risk of atherosclerosis and resultant cardiovascular complications ^3^. Although the hepatic vascular endothelium is reported to have unique barrier functions relevant to lipid metabolism and transport, it is unclear whether this tissue affects lipid transport and impacts atherosclerosis and hepatic steatosis ^4–6^.

Microsomal triglyceride transfer protein (MTTP) plays an important role in hepatic and intestinal lipid metabolism and transport. MTTP is indispensable for the assembly and secretion of very-low-density lipoprotein (VLDL) particles from hepatocytes by transferring triglycerides and phospholipids to nascent apolipoprotein B (apoB) ^7,8^. Disruption of MTTP leads to impaired VLDL secretion, hepatic steatosis, and altered circulating plasma lipid levels ^7,9^. Importantly, MTTP function has also been linked to atheroprogression, as changes in MTTP activity can influence plasma lipid profiles and the development of atherosclerosis ^10,11^.

Biosynthesis of phospholipids is catalyzed by choline ethanolamine phosphotransferase 1 (CEPT1). It is responsible for the production of the majority of phosphatidylcholine (PC) and phosphatidylethanolamine (PE), which are essential phospholipids needed for cell membrane structure and intracellular signaling (12,13). CEPT1-derived phospholipid derivatives also activate peroxisome proliferator-activated receptor α (PPARα), a nuclear receptor transcription factor that induces fatty acid β-oxidation ^14^. Conditional knockdown of *Cept1* in the endothelium results in reduced endothelial cell (EC) function and signaling, as well as reduced post-ischemia tissue recovery ^13,14^. Interestingly, in this setting post-ischemia tissue recovery is rescued with the administration of a PPARα agonist, demonstrating that CEPT1 may be an important master regulator of EC health and function. Notably, PPARα activation is known to upregulate MTTP expression, linking lipid metabolism and vascular recovery through coordinated signaling ^15^.

Direct interactions between CEPT1 and MTTP have not been demonstrated. Given their parallel signaling via PPARα and overlapping function in lipid metabolism and homeostasis, we hypothesized that endothelial CEPT1 may influence hepatic MTTP to regulate circulating lipids, alter the course of hepatic steatosis, and impact atheroprogression.

## Materials and Methods

### Data availability statement

The data that supports the findings of this study are available from the corresponding author upon reasonable request.

### Human liver tissue collection and ethical approval

Human liver tissue samples were obtained from patients at a single university-affiliated medical center undergoing clinically indicated liver biopsies or hepatic resections. Specimens were classified as steatotic or non-steatotic based on histopathological assessment. Acquisition and use of biobanked hepatic tissue is approved by the Human Research Protection Office (HRPO #201602080). Only excess tissue not required for clinical diagnosis was used for research purposes and testing of these specimens were conducted in compliance with institutional ethical guidelines. Liver tissue specimens were used for Western blot analysis, histological staining, and assessment of MTTP activity.

### Transfection and *in vitro* co-culturing

Human umbilical vein endothelial cells (HUVECs) were seeded at a density of 1 × 10^6^ cells per well in 6-well plates. Cells were then serum-starved for 2 hours in EBM2 medium. Transfection was performed using Lipofectamine™ RNAiMAX transfection reagent (ThermoFisher, No.13778075) and either *Cept1* siRNA (EHU064851-50UG, Mission eSIRNA Human *Cept1*) or *Pparα* siRNA (ThermoFisher, Cat#142804,10 µM) following the manufacturer’s protocol. After 6 hours, the transfection medium was replaced with complete EGM media. The following day HepG2 cells were added to the transfected HUVECs or controls for co-culture experiments. Fenofibrate (FEN), a PPARα agonist, was added to the co-culture at a final concentration of 50 µM. The co-cultures were incubated for 48 hours at 37°C in 5% CO_2_. RNA and protein were extracted from triplicate wells using Trizol reagent and RIPA buffer, respectively, for subsequent analyses.

### MTTP activity assay

MTTP activity was measured using the Sigma Aldrich ROAR MAK110 kit, following the manufacturer instructions. Fluorescence was measured using a fluorescence spectrometer (λ excitation = 465 nm, λ emission = 535 nm) after equilibrating the plate at 37°C. Control wells contained 198 µL of MTTP assay buffer, 4 µL of donor particles, and 4 µL of acceptor particles, while experimental wells included additional samples: HepG2 cell lysates or mice liver homogenates and human steatotic/non-steatotic liver samples, each standardized to 10-25 µg of protein per 200 µL reaction. All samples were prepared in triplicate. Fluorescence readings were recorded every 15 minutes for 3-4 hours until the fluorescence signal stabilized. MTTP activity was then determined and normalized to the protein concentration in each sample, allowing to ensure an accurate comparison of enzyme activity across different samples.

### Real-time PCR

Hepatic ECs were isolated from mouse livers using positive cell sorting with PECAM-1- and/or ICAM-2-coated magnetic beads, as previously described.(11) RNA was extracted and purified from ECs or hepatic tissue, HUVEC, and HepG2 cells and converted to cDNA. For whole liver RNA extraction, liver tissue was homogenized, and total RNA was isolated using established protocols. Quantitative PCR was performed using SYBR Green Master Mix (Applied Biosystems). The threshold cycle (Ct) values were normalized to *CEPT1*, *MTTP*, and *PPARα* mRNA and expressed as fold change relative to the mean Ct value of control using the ΔCt method.

### Animals, diets, and treatments

All experimental procedures, including housing, breeding, blood and tissue collection, diets, and treatments, were conducted by national guidelines and regulations and approved by the institutional animal care and use committee (IACUC). Wild-type (*C57BL/6J*) mice and *Apoe^-/-^*mice of both sexes were obtained from Jackson Laboratory. Conditional endothelial *Cept1* knockdown was achieved using a tamoxifen-inducible (0.05mg/g of body weight administrated intraperitoneal for five consecutive days) *Cre-loxP* system, with *loxP* sites flanking exon 3 of the *Cept1* gene. (11) To generate *Cept1^fl/fl^Cre^+^Apoe^-/-^*mice, we crossed *Cept1^fl/fl^Cre^+^* mice with *Apoe^-/-^* mice. Offspring were genotyped by PCR analysis of tail DNA to confirm the presence of floxed *Cept1* alleles, Cre recombinase, and the *Apoe^-/-^* allele, following protocols utilizing loxP and Cre-mediated recombination as described in established methods ^16^. At 6 weeks of age, mice of both sexes were randomly assigned to experimental groups and placed on a Western-type diet (HFD, 42% fat content) for 12 weeks ^17^. To induce *Cre* expression and *Cept1* knockdown, mice received daily intraperitoneal tamoxifen injections (50 mg/kg) for 5 consecutive days starting at 8 weeks of age. Body weight was measured weekly throughout the high-fat diet period, and blood samples were collected every 4 weeks.

### Serum, liver and stool lipidomic and plasma measurements

Cholesterol and triglyceride concentrations in the serum, liver homogenate and stool of mice were assayed using commercially available kits (Triglyceride Infinity Kit TR22421 and Cholesterol Infinity Kit TR13421, Thermo Scientific). To measure serum and stool-free fatty acids (FFA), an enzymatic colorimetric assay (NEFA-HR2, Fujifilm Wako Diagnostics) was used according to the manufacturer instructions.

### Immunohistochemistry

For immunofluorescence staining, freshly harvested aortic valve and liver tissue from euthanized mice, as well as human liver tissues (as described previously), were first incubated in 30% fructose for 2 hours at room temperature, followed by 15% fructose overnight. The next day, tissues were embedded in the OCT compound and frozen at -20°C. Sections were then cut at a thickness of 10 µm. They were blocked with 10% donkey serum for 1 hour at room temperature. Sections were incubated overnight at 4°C with the following primary antibody anti-CD68 (1:200, Bio-Rad, MCA17576A), CD31 (1:250, Santa Cruz, sc-376764), MTTP (1:150, Thermo Fischer, PA5-76049) and CEPT1 (1:250, Bioss, bs-12284R-Cy3). After washing the slides in PBS, slides were incubated with Alexa Fluor-488 donkey anti-rat (1:500; Jackson ImmunoResearch, West Grove, USA) secondary antibody for 1 hour at room temperature, followed by washes. Images were acquired using a Leica Thunder DM6 B Microsystems inverted fluorescence microscope. CD68, CD31 & CEPT1 stained cells were analyzed by applying color thresholds. For histological analysis, adjacent sections were stained with hematoxylin and eosin (H&E). For all histological and imaging endpoints, researchers quantifying data were blinded to experimental group allocation throughout analysis.

### Aortic valve histology and lesion assessment

Using magnified dissection mouse hearts were excised *en bloc* and embedded in OCT compound. The hearts were oriented to facilitate cross-sectioning at the aortic annulus using a cryostat in 10 µm thickness sections. Cross sections of the aortic annulus were then stained with 0.5% (w/v) ORO for 10 minutes as well as H&E, and subsequently imaged. The ORO-positive intimal lesion surface area was measured and expressed as μm² or as a percentage of the total intimal area. The quantification was performed by an observer blinded to experimental group.

### Western blot analysis

GAPDH, MTTP, and CEPT1 protein levels, were evaluated using Western blotting. Cell and liver tissue samples were lysed in RIPA buffer supplemented with protease inhibitors, followed by homogenization and centrifugation to obtain clarified lysates. Protein concentrations were determined using a Bradford assay. Equal amounts of protein (25 µg) were mixed with 2X SDS loading buffer, heated, and separated on a 10% SDS-PAGE gel. Proteins were then transferred to PVDF membranes. Membranes were blocked with 5% BSA in TBS-T and incubated overnight at 4°C with primary antibodies: GAPDH (1:1,000, Cell signaling, 2118), MTTP (1:1,500, ThermoFischer, PA5-76049), and CEPT1 (1:2000Bioss, bs-12284R-Cy3). Following washes, membranes were incubated with HRP-conjugated secondary antibodies (1:2,000). Protein bands were visualized using chemiluminescence and captured using a digital imaging system. Band intensities were normalized to GAPDH as a loading control, and relative protein expression was quantified using image analysis software.

### Aortic effacement and atherosclerotic plaque assessment

Following euthanasia, the aorta was perfused and flushed with PBS saline via left ventricular puncture. The entire aorta, extending to the iliac bifurcation, was carefully excised under magnified tissue dissection and fixed in 4% paraformaldehyde for 24 h. Subsequently, the aorta was longitudinally opened and pinned in a square dish on a black wax surface. The prepared aorta was stained with Oil Red O (ORO, 0.5% (w/v)) for visualization and quantification of atherosclerotic lesions. Fixed samples were rinsed with 60% isopropanol for 30 seconds, then incubated in ORO working solution for 15–30 minutes. After staining, samples were rinsed briefly with 60% isopropanol, then with deionized water for 1 minute, and counterstained with Mayer’s hematoxylin for 1 minute. Slides were mounted with Crystal/Mount and sealed with nail polish before imaging. After staining, high-resolution photographs were taken for further analysis. En-face quantification of atherosclerotic lesions was performed using ImageJ software.

### Experimental design, randomization, and blinding

Mice and cell culture samples/wells were randomly assigned to experimental groups to reduce selection bias. Randomization was applied during sample allocation and, where possible, during data acquisition. Experimental conditions and sample processing were organized to minimize confounding factors. Researchers performing experiments and analyses were not blinded to group allocation; however, outcome measurements that required subjective assessment (e.g., histological scoring) were performed using pre-established criteria.

### Statistical analysis

Data were expressed as mean ± SEM or percentage. For comparing two groups, either the student’s t-test or the non-parametric Mann-Whitney U test was used for continuous variables. The difference between groups at different time points was analyzed using two-way ANOVA with repeated measures. A *p*<0.05 was considered statistically significant. All analyses were performed using GraphPad Prism.

## Results

### Human hepatic steatosis is associated with altered CEPT1 and MTTP

We studied samples obtained from patients with and without hepatic steatosis. The two patient groups had similar sex distribution, diabetes prevalence, and alcohol consumption; however, BMI was higher in the liver steatosis group compared to the non-steatosis group (Table 1). Analysis of human liver samples revealed interesting differences between non-steatotic (n=11) and steatotic (n=12) conditions (Fig.1). For each human liver sample, MTTP was evaluated using five replicate measurements, and demonstrated significantly reduced enzyme activity in steatotic human liver tissue (Fig.1A; p<0.01). Total levels of *CEPT1* and *MTTP* mRNA were not significantly different between steaototic and non-steaototic liver tissue (Supplementary Fig.1). Western blot analysis demonstrated no significant difference in CEPT1, PPARα or MTTP content in steatotic and non-steaotitic tissue (Fig.1B-D; p<0.05). Immunofluorescence analysis also showed a modest decrease of MTTP (Fig. 1E-G), and a notable reduction of CEPT1:CD31 co-localization in steatotic liver sections (Fig.1H; p<0.01).

**Figure 1:**
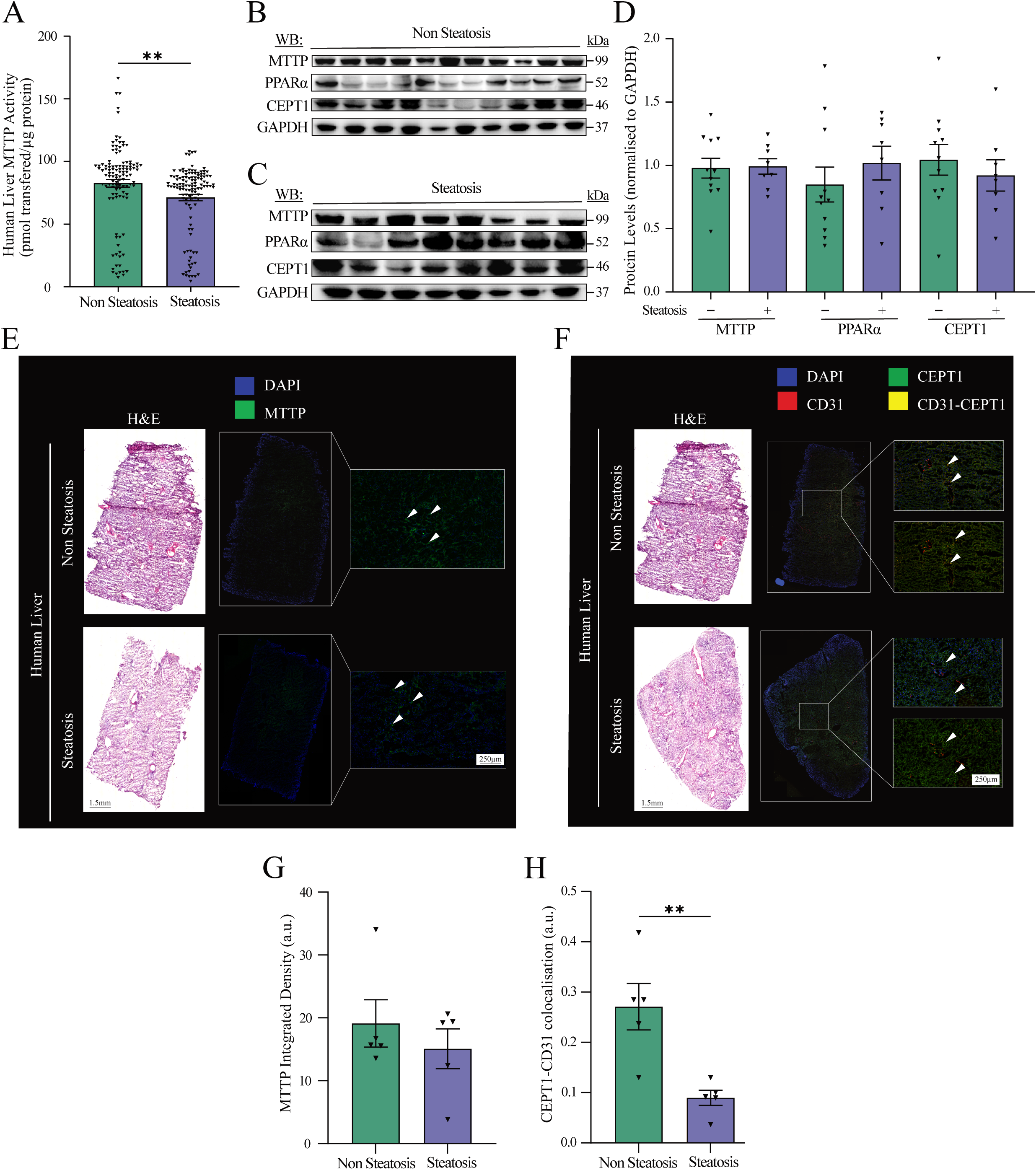
Endothelial CEPT1 regulates MTTP in human liver steatosis. (A) Quantification of human MTTP activity at the plateau phase (170-210 min) non-steatosis (n=11) and steatosis samples (n=12). Each patient is represented by a group of approximately 5 data points (dots). (B-C) Representative western blot showing reduced CEPT1 and MTTP protein levels in human liver tissue from non-steatosis (n=11) and steatosis samples (n=12). GAPDH serves as a loading control. (D) Quantification of hepatic CEPT1 (C) and MTTP (D) protein levels. (E-F) Representative immunofluorescence images of human liver sections, comparing non-steatosis and steatosis conditions: (E) H&E staining with DAPI (blue) and MTTP (green); white arrows highlight regions of positive MTTP staining; (F) H&E staining alongside DAPI (blue), CD31 (red), and CEPT1 (green), white arrows indicate areas of CD31–CEPT1 colocalization, shown in yellow; (G) Quantification of MTTP integrated density, showing a significant decrease in steatosis (*p<0.05). (H) Quantification of CEPT1/CD31 co-localization, which is significantly reduced in steatotic samples (**p<0.01). Data represent mean ± SEM. Statistical significance was determined by Unpaired t-test.

**Table 1.**
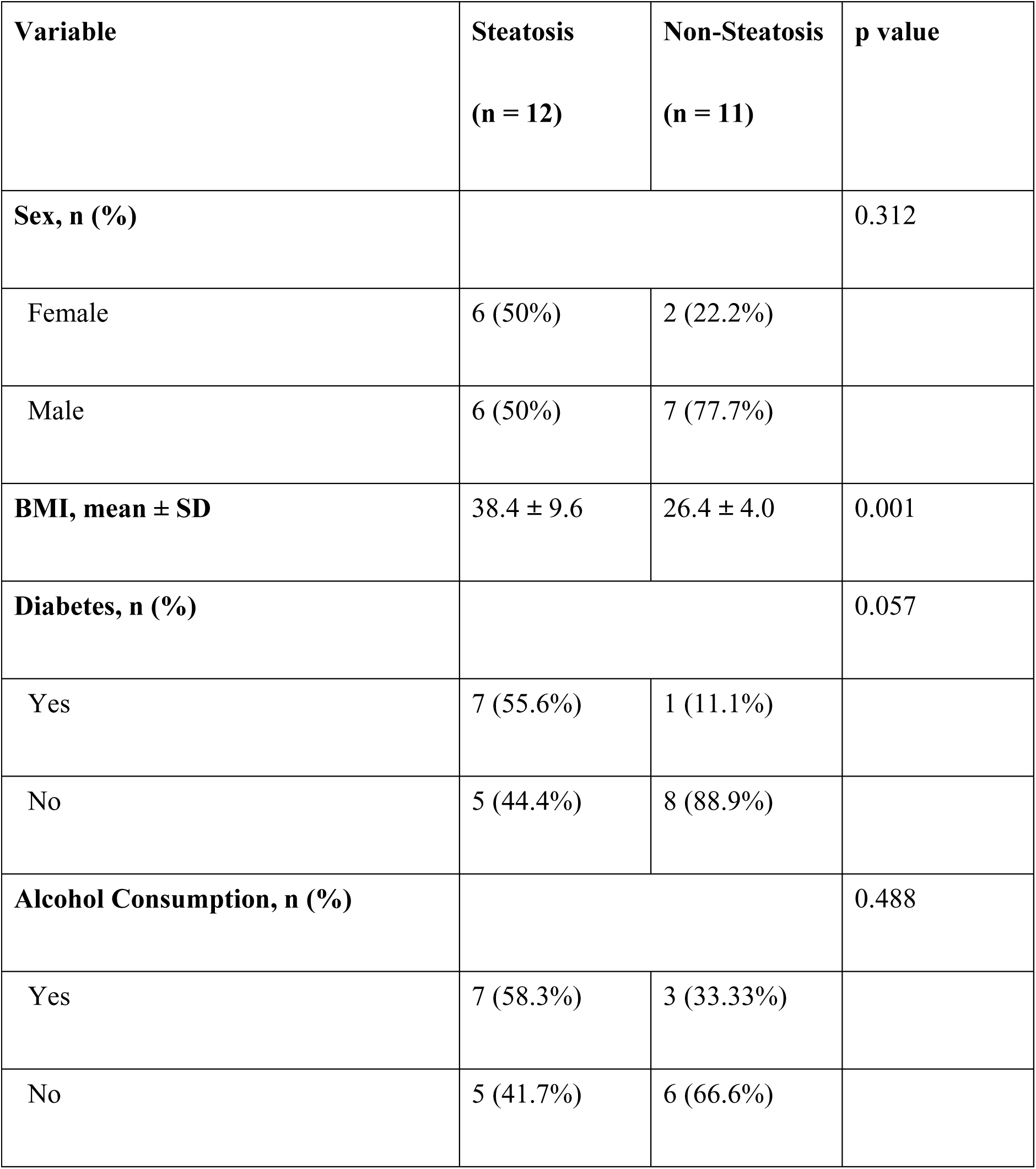
Patient Demographic and Clinical Characteristics.

### Endothelial CEPT1 modulates hepatic MTTP via PPARα signaling

Based on our findings of reduced MTTP activity in human steatotic liver samples, we evaluated whether CEPT1 deficiency in ECs can directly impact hepatic cell MTTP activity *in vitro* (Fig.2A). HUVECs transfected with *CEPT1* siRNA (si*CEPT1*+) were co-cultured with HepG2 cells (Fig. 2B). HUVECs si*CEPT1*+ had reduced *CEPT1* (Fig.2C; p<0.0001) and *PPARα* (Fig.2D; p=0.008), but no change in *MTTP* (Fig. 2E). Interestingly, HUVECs with *CEPT1* knockdown led to decreased *MTTP* (Fig.2F; p=0.02) and *PPARα* (Fig.2G; p<0.0001) in co-cultured HepG2 cells. Additionally, there was reduced HepG2 MTTP activity (Fig.2H-I; p=0.03) compared to other co-culture groups. These results suggest that endothelial *CEPT1* leads to paracrine regulation of hepatic cellular *MTTP* expression and MTTP activity.

**Figure 2:**
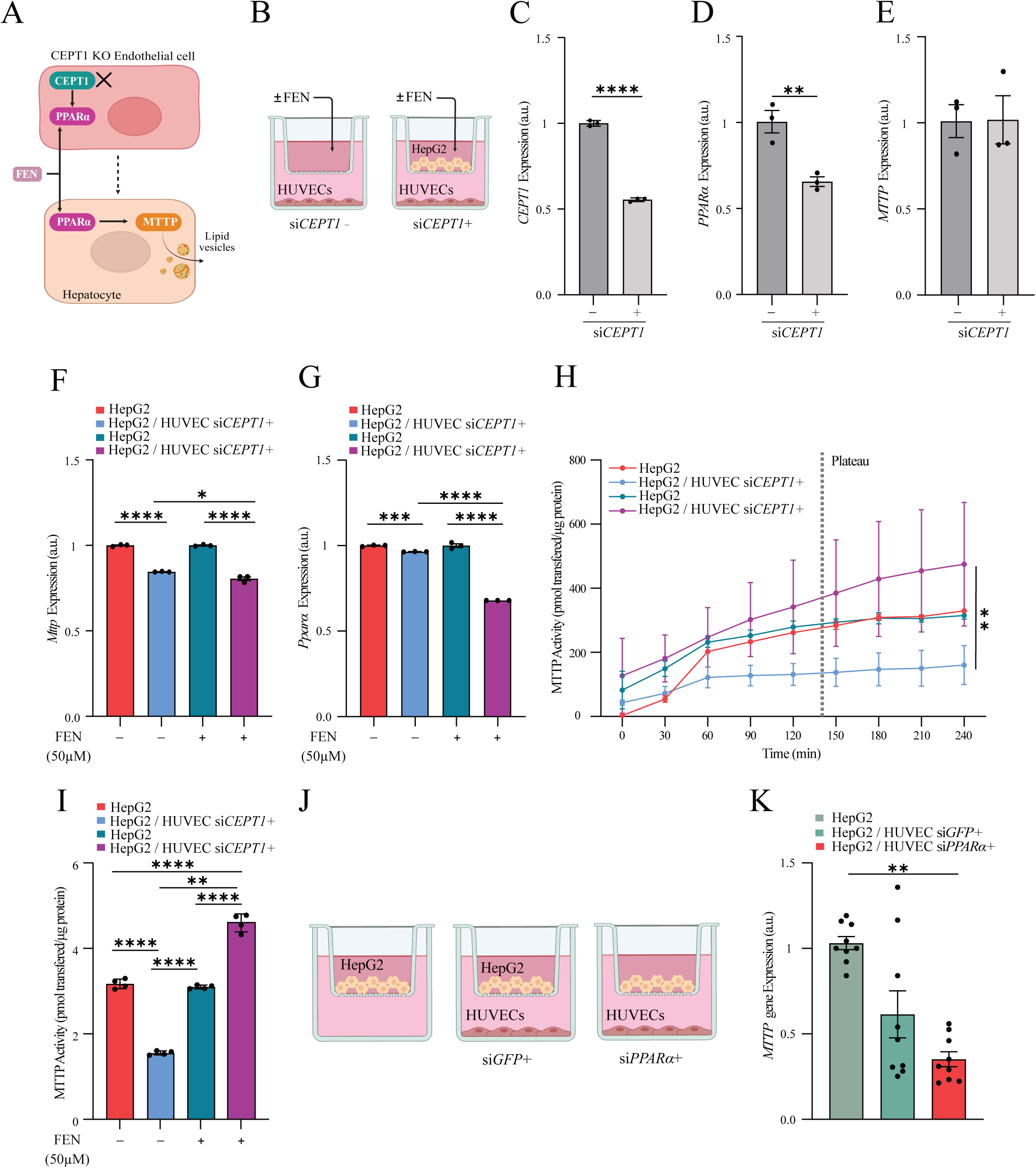
*In vitro CEPT1* knockdown in HUVECs alters MTTP in HepG2 cells via PPARα. (A) Schematic illustrating the proposed impact of endothelial CEPT1 (EC CEPT1) on hepatic MTTP and PPARα regulation. EC CEPT1 influences hepatocyte function, potentially impacting VLDL assembly and fatty acid metabolism. Fenofibrate (FEN), a PPARα activator, is included to indicate pharmacological activation of PPARα in this pathway (B) Experimental setup illustrating co-culture of HUVECs with HepG2 cells ± FEN treatment (50 µM) for 48h. HUVECs were transfected with si*CEPT1*+. (C-E) Relative mRNA expression of *CEPT1* (C), *PPARα* (D), and *MTTP* (E) in HUVECs transfected with siRNA targeting *Cept1* compared to untransfected HUVECs (control), normalized to controls (n=3). Relative mRNA expression levels of *MTTP* (F) and *PPARα* (G) in HepG2 cells co-cultured with HUVECs ± siRNA targeting *CEPT1* and ±FEN treatment (50µM) for 48h, normalized to controls (n=3). (H) MTTP activity curve (pmol transferred/µg protein) over time for HepG2 cells from control and co-culture with HUVEC si*CEPT1*+ cells ±FEN treatment (50µM for 48h). Plateau indicates maximal MTTP transfer capacity, as determined by endpoint assay guidelines. (I) Quantification of MTTP activity at the plateau phase (195-240 min) for HepG2 cells from control and co-culture with HUVEC si*CEPT1*+ cells ±FEN treatment. Data were analyzed by Mann-Whitney U test; n=4. (J) Experimental setup illustrating co-culture of HepG2 with si*PPARα*+ transfected HUVECs vs control (HUVECs si*PPARα*-) for 48h. (K) Relative mRNA expression of *MTTP* in HepG2 co-culture with HUVECs si*PPARα*+ vs positive control (HUVECs si*GFP*+) and negative control (un-transfected HUVECs) (n=9). Data were analyzed by t-test or one-way ANOVA followed by Tukey’s multiple comparisons test. *p<0.05, **p<0.01, ***p<0.001, ****p<0.0001.

To investigate whether PPARα activation could rescue the reduction in MTTP activity caused by endothelial *siCEPT1*+, we treated co-cultured HepG2 cells with the PPARα agonist fenofibrate (FEN). FEN treatment successfully restored MTTP activity in HepG2 cells that were co-cultured with HUVECs si*CEPT1*+ (Fig.2H-I; p=0.006). This suggested that PPARα activation can compensate in HepG2 cells for the impact of selective knockdown of *CEPT1* in co-cultured ECs. Conversely, siRNA-mediated knockdown of *PPARα* in HUVECs (HUVEC si*PPARα+)* co-cultured with HepG2 cells resulted in a significant reduction in HepG2 *MTTP* expression (Fig.2J-K; p=0.001) compared to control (non-transfected HUVECs), further demonstrating that PPARα was likely acting downstream of CEPT1 in ECs and was contributing to the paracrine regulation of MTTP in HepG2 cells.

### Conditional knockdown of *Cept1* impacts murine serum lipids

We next explored how conditional knockdown of *Cept1* in the endothelium impacts lipid metabolism and transport *in vivo*. Four murine experimental groups were utilized: *Cept1^fl/fl^Cre^+^Apoe^-/-^*(n=15), *Cept1^fl/fl^Cre^-^Apoe^-/-^* (n=10), *Cept1^fl/fl^Cre^+^Apoe^+/+^* (n=17), and *Cept1^fl/fl^Cre^-^Apoe^+/+^*(n=9; Fig.3A). All groups were maintained on HFD for at least 12 weeks. Compared to controls, *Cept1^fl/fl^Cre^+^Apoe^+/+^*mice demonstrated no significant difference in body weight (Fig.3B), no difference in circulating serum cholesterol (Fig.3C), significantly reduced serum TG (Fig.3D; p<0.0001), and no change in free fatty acids (FFA; Fig.3E). *Cept1^fl/fl^Cre^+^Apoe^-/-^*had a notable reduction in circulating serum cholesterol (Fig.3C; p=0.013), triglycerides (TG; Fig.3D; p=0.003), and free fatty acid (FFA; Fig.3E; p=0.004). These findings align with previous reports linking endothelial phospholipid metabolism to systemic lipid regulation ^14^ and show that endothelial *Cept1* plays an important role in regulating circulating serum lipids.

**Figure 3.**
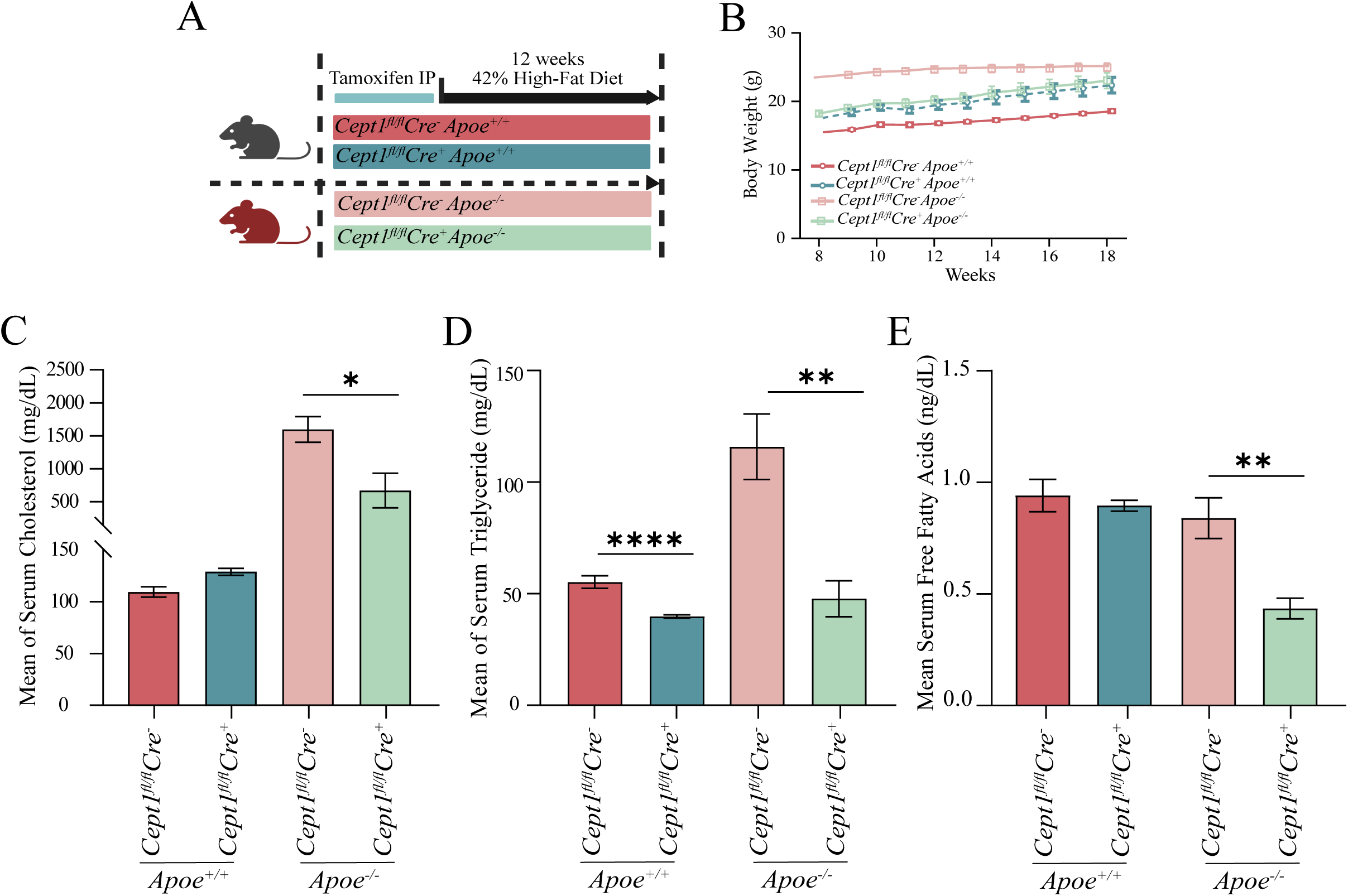
Impact of endothelial *Cept1* knockdown on serum lipid profiles *in vivo.* (A) Schematic representation of experimental design. Tamoxifen was administered intraperitoneally to induce Cre recombinase activity in *Cept1^fl/fl^Apoe^+/+^* and *Cep1^fl/fl^ Apoe^-/-^* mice. After tamoxifen treatment, mice were fed a 42% high-fat diet for 12 weeks. Serum samples were collected weekly for further analysis. (B) Body weight measurements (in grams) of mice over 12-week period. (C-E) Serum lipid profile after 12 weeks on a high-fat diet. Mean serum levels of total cholesterol (mg/dL) (C), triglycerides (mg/dL) (D), and free fatty acids (ng/dL) (E) in *Cept1^fl/fl^Apoe^+/+^* and *Cep1^fl/fl^ Apoe^-/-^* mice compared to controls. All data are presented as mean ± SEM. Statistical significance was determined by Unpaired t-test. *p<0.05, **p<0.01, ****p<0.0001, and n ≥ 6 for all groups.

To elucidate whether reduced systemic circulating lipids are confounded by intestinal malabsorption of circulating lipid, we also evaluated fecal lipid content in all murine groups. Compared to *Cept1^fl/fl^Cre^-^Apoe^+/+^* mice, *Cept1^fl/fl^Cre^+^Apoe^+/+^* mice demonstrated no statistically significant differences in fecal total cholesterol (Supplementary Fig.2A), triglycerides (Supplementary Fig.2B), and FFAs (Supplementary Fig.2C). Compared to *Cept1^fl/fl^Cre^-^Apoe^-/-^* mice, *Cept1^fl/fl^Cre^+^Apoe^-/-^*mice also demonstrated no significant difference in fecal total cholesterol, triglycerides, and FFA (Supplementary Fig.2D-F). These results suggest that the reduction in serum lipids of mice with conditional endothelial *Cept1* knockdown (*Cept1^fl/fl^Cre^+^*) is not due to an underlying intestinal lipid malabsorption phenotype.

### EC-specific *Cept1* knockdown impairs hepatic lipid metabolism via downregulation of MTTP

Livers from all murine groups were also evaluated using ORO staining to evaluate hepatic lipid content. Lipid content in *Cept1^fl/fl^Cre^+^Apoe^+/+^*mice showed significantly reduced hepatic lipid droplets compared to *Cept1^fl/fl^Cre^-^Apoe^+/+-^*mice (Fig.4A&B; p=0.004). However, the lipid content was unchanged between *Cept1^fl/fl^Cre^-^Apoe^-/-^*and *Cept1^fl/fl^Cre^+^Apoe^-/-^* mice (Fig.4A&B). We also observed no difference in hepatic content of triglycerides, total cholesterol, and FFAs among all mouse groups (Supplementary Fig.3A-C).

**Figure 4.**
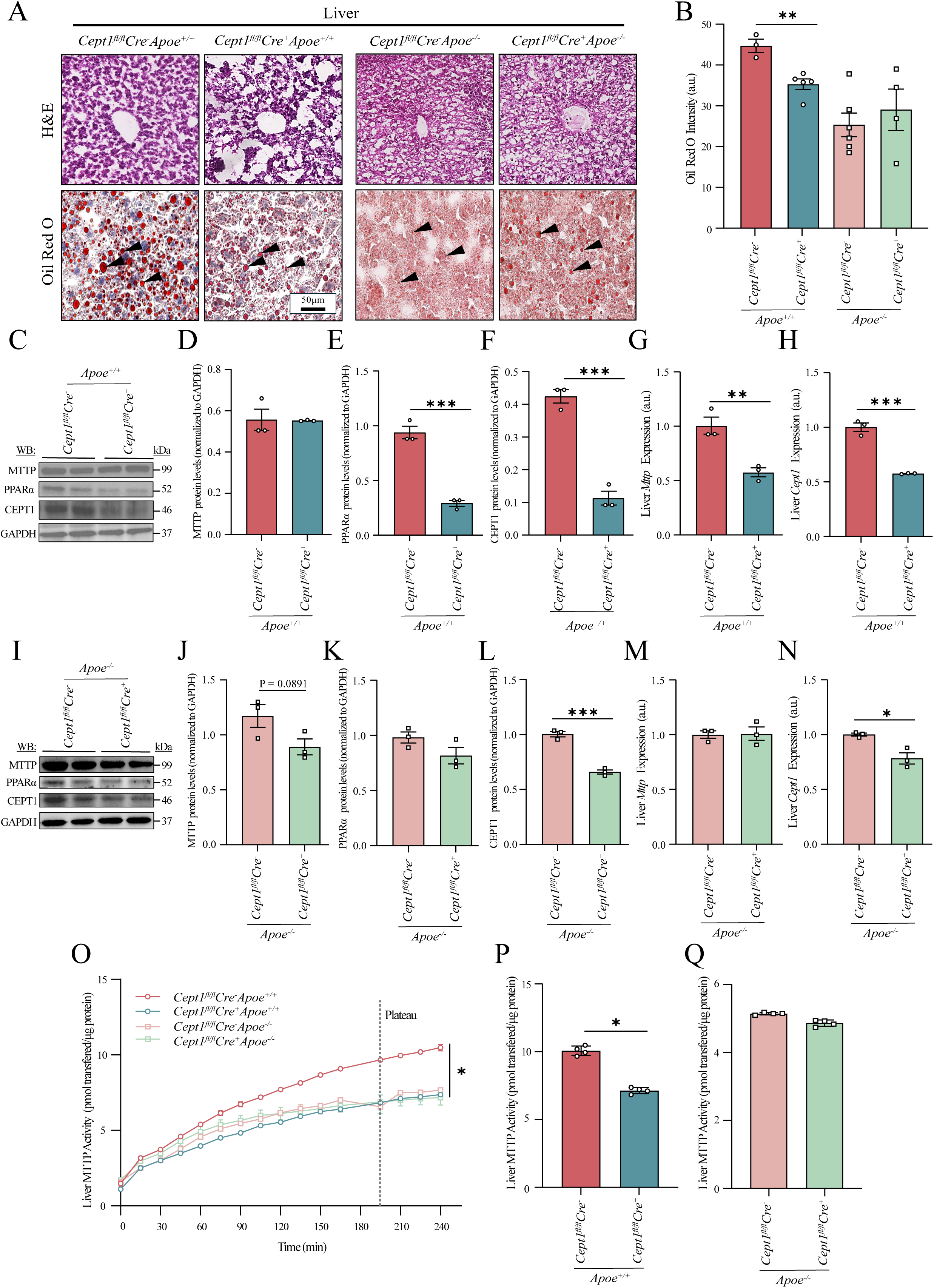
EC *Cept1* knockdown alters lipid metabolism and hepatic gene expression *in vivo.* (A) Representative H&E-stained liver sections from *Cept1^fl/fl^Cre^+^Apoe^-/-^, Cept1^fl/fl^Cre^-^Apoe^-/-^*, *Cept1^fl/fl^Cre^+^Apoe^+/+^*and *Cept1^fl/fl^Cre^-^Apoe^+/+^* mice. Scale bar = 100 μm. (B) Quantification of hepatic lipid accumulation as measured by ORO intensity in liver sections across all experimental groups. Data analyzed by one-way ANOVA with Tukey’s post-hoc test; n=4-6 per group; **p<0.01. (C) Representative Western blots of MTTP, PPARα, CEPT1, and GAPDH in liver lysates of *Cept1^fl/fl^Cre^-^Apoe^+/+^*and *Cept1^fl/fl^Cre^+^Apoe^+/+^*. Molecular weights (kDa) are indicated. Quantification of hepatic *Cept1^fl/fl^Cre^-^Apoe^+/+^ and Cept1^fl/fl^Cre^+^Apoe^+/+^* MTTP (D), PPARα (E), CEPT1 (F) protein levels normalized to GAPDH in all mice groups; n=3. Hepatic *Mttp* (G) and *Cept1* (H) gene expression in *Cept1^fl/fl^Cre^-^Apoe^+/+^* and *Cept1^fl/fl^Cre^+^Apoe^+/+^*. (I) Representative Western blots of MTTP, PPARα, CEPT1, and GAPDH in liver lysates of *Cept1^fl/fl^Cre^-^Apoe^-/-^* and *Cept1^fl/fl^Cre^+^Apoe^-/-^*. Molecular weights (kDa) are indicated. Quantification of hepatic *Cept1^fl/fl^Cre^-^Apoe^+/+^*and *Cept1^fl/fl^Cre^+^Apoe^-+/+^* MTTP (J), PPARα (K), CEPT1 (L) protein levels normalized to GAPDH in all mice groups; n=3. Hepatic *Mttp* (M) and *Cept1* (N) gene expression in *Cept1^fl/fl^Cre^-^Apoe^-/-^* and *Cept1^fl/fl^Cre^+^Apoe^-/-^*. Data were analyzed by t-test. (O) Liver MTTP activity curve (pmol transfered/μg protein) over time in *Cept1^fl/fl^Cre^+^Apoe^-/-^,* and *Cept1^fl/fl^Cre^+^Apoe^+/+^*mice compared to controls. Plateau represents the maximal transfer capacity achieved under these experimental conditions, as per manufacturer’s guidelines for endpoint assays. (P-Q) MTTP activity at the plateau phase (180-240 min) in *Cept1^fl/fl^Cre^+^Apoe^+/+^* (P), and *Cept1^fl/fl^Cre^+^Apoe^-/-^* (Q) mice compared to controls. Data were analyzed by Mann-Whitney U test; n=4. *p<0.05, **p<0.01, ***p<0.001, ****p<0.0001.

Immunoblot analysis and quantification showed that EC-specific deletion of *Cept1* in *Cept1^fl/fl^Cre^+^Apoe^+/+^* resulted in no change in hepatic MTTP content compared to controls (Fig.4C&D). However, there was a significant reduction in hepatic PPARα (Fig.4E; p=0.001) and CEPT1 (Fig.4F; p=0.001). Similarly, *Cept1^fl/fl^Cre^+^Apoe^+/+^* mice had decreased hepatic *Mttp* (Fig.4G, p=0.008) and *Cept1* (Fig.4H; p=0.0004) expressions*. Cept1^fl/fl^Cre^+^Apoe^-/-^*also had no change in hepatic MTTP and PPARα content (Fig.4I-K) but decreased CEPT1 (Fig.4L). At the gene level, no difference was observed in hepatic *Mttp* (Fig.4M), but *Cept1* was significantly decreased (Fig.4N).

Dynamic functional assessment of hepatic MTTP showed reduced enzymatic activity in *Cept1^fl/fl^Cre^+^Apoe^+/+^*, *Cept1^fl/fl^Cre^-^Apoe^-/-^*, *Cept1^fl/fl^Cre^+^Apoe^-/-^*mice, relative to *Cept1^fl/fl^Cre^-^ Apoe^+/+^* mice (Fig.4O). Endothelial knockdown of *Cept1* alone demonstrated a significant impact on hepatic MTTP activity (Fig.4P; p=0.028), but this difference was no longer observed in *Apoe^-/-^* mice (Fig.4Q). Collectively, these findings indicate that while knockdown of endothelial *Cept1* can clearly impact hepatic *Mttp* expression (Fig.4G) and MTTP activity (Fig.4O&P), these observations are specific to *Apoe^+/+^* mice. This suggests that severe hyperlipidemia in *Apoe^-/-^* mice is an independent factor that may overcoming the impact of CEPT1 on MTTP.

### Conditional knockdown of *Cept1* in ECs decreases atherosclerosis

Since reduced hepatic MTTP activity can lead to reduced lipid export from the liver and circulating serum lipids, we also sought to evaluate the impact of conditional knockdown of endothelial *Cept1* on aortic atherosclerotic plaque formation. Compared to *Cept1^fl/fl^Cre^-^Apoe^+/+^*(n ≥ 5), *Cept1^fl/fl^Cre^+^Apoe^+/+^* (n ≥ 5) mice demonstrated markedly reduced aortic plaque (Fig.5A-E, p=0.006). Similarly, compared to *Cept1^fl/fl^Cre^-^Apoe^-/-^* (n ≥ 5) mice, *Cept1^fl/fl^Cre^+^Apoe^-/-^* (n ≥ 5) mice demonstrated significantly reduced total aortic plaque (Fig.5F-G; p=0.008, and Supplementary Fig. 4A), including in the aortic arch (Fig.5H, p=0.046), thoracic aorta (Fig.5I; p=0.016), and aortoiliac segments (Fig.5J; p<0.01). These findings suggest that endothelial *Cept1* plays a vital role in aortic plaque formation. Further analysis of lipid deposition in the aortic annulus showed significantly reduced ORO-stained atherosclerotic lesions in both *Cept1^fl/fl^Cre^+^Apoe^+/+^*and *Cept1^fl/fl^Cre^+^Apoe^-/-^* (Fig.5K-M; p=0.0007 and p=0.02, respectively). CD68+ macrophage content in the aortic annulus was also notably reduced in *Cept1^fl/fl^Cre^+^Apoe^-/-^* (Fig.5N&O; p=0.04).

**Figure 5.**
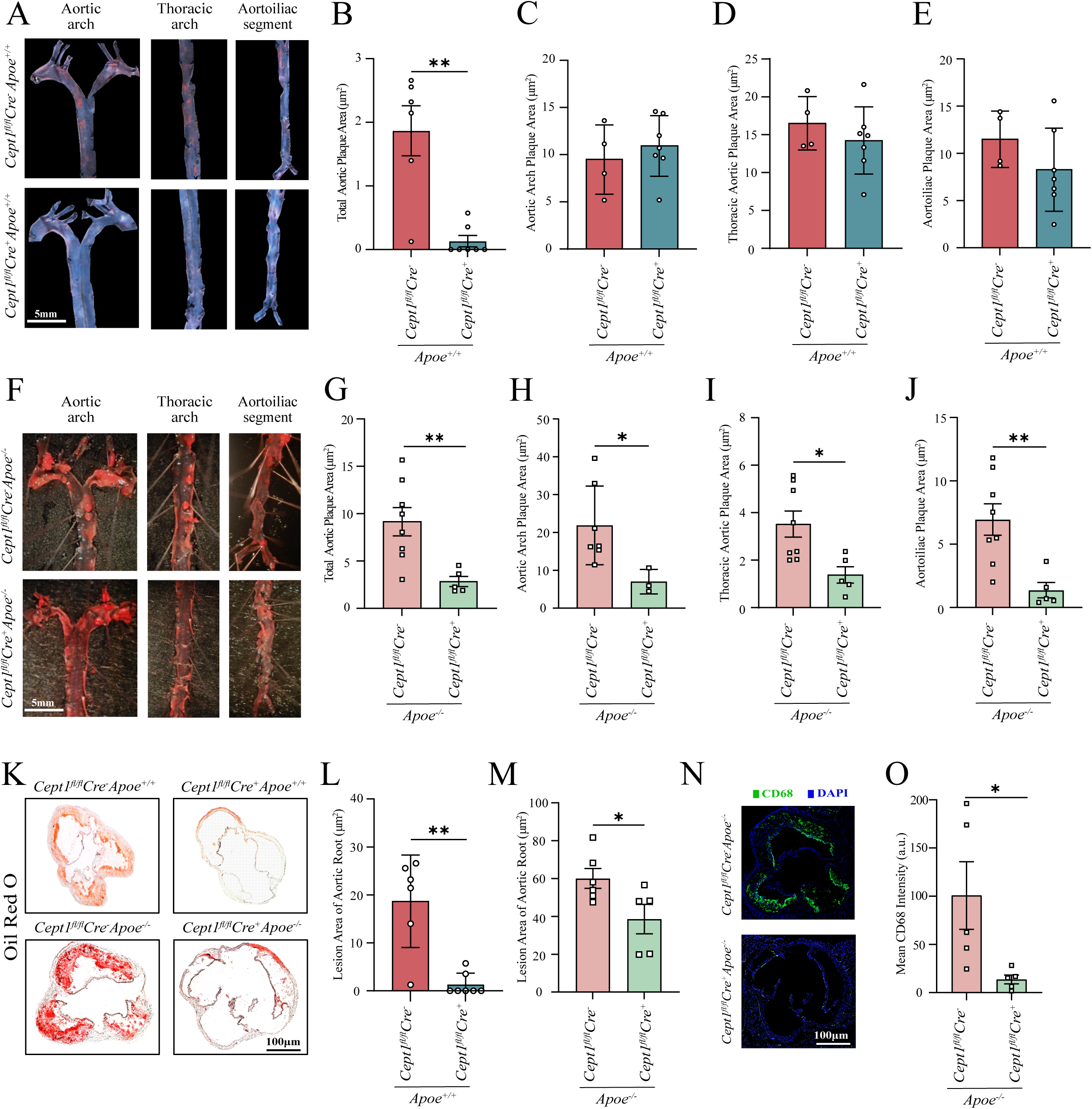
EC *Cept1* knockdown reduces atherosclerotic plaque burden *in vivo.* Representative images of ORO stained *en face* preparations of the aortas in *Cept1^fl/fl^Cre^-^Apoe^+/+^* and *Cept1^fl/fl^Cre^+^Apoe^+/+^*(A). Scale bar=5mm. Quantification of plaque area in the total aorta (B; n ≥ 5), aortic arch (C; n ≥ 4), thoracic aorta (D; n ≥ 5), and aortoiliac segment (E; n ≥ 4) of *Cept1^fl/fl^Cre^-^Apoe^+/+^,* and *Cept1^fl/fl^Cre^+^Apoe^+/+^* mice. Representative images of ORO stained *en face* preparations of the aortas in *Cept1^fl/fl^Cre^-^Apoe^-/-^*and *Cept1^fl/fl^Cre^+^Apoe^-/-^* (F). Scale bar=5mm. Quantification of plaque area in the total aorta (G; n ≥ 5), aortic arch (H; n ≥ 4), thoracic aorta (I; n ≥ 5), and aortoiliac segment (J; n ≥ 4) of *Cept1^fl/fl^Cre^-^Apoe^-/-^,* and *Cept1^fl/fl^Cre^+^Apoe^-/-^*mice. (K) Representative images of ORO-stained aortic root sections from *Cept1^fl/fl^Cre^+^Apoe^+/+^* and *Cept1^fl/fl^Cre^+^Apoe^-/-^* mice compared to controls. Scale bar=100µm. (L-M) Quantification of lipid deposition in the aortic root sections; n ≥5 *Cept1^fl/fl^Cre^+^Apoe^+/+^*(L) and *Cept1^fl/fl^Cre^+^Apoe^-/-^* (M) compared to their corresponding controls. (N) Representative immunofluorescence staining for CD68 (green) and DAPI (blue) in the aortic root of *Cept1^fl/fl^Cre^-^Apoe^-/-^* and *Cept1^fl/fl^Cre^+^Apoe^-/-^* mice. Scale bar=100μm. (O) Quantifying means CD68 intensity (arbitrary units, a.u.) (n=5). t-tests determined statistical significance. *p<0.05, **p<0.01.

## Discussion

Our findings reveal that circulating serum lipids and atheroprogression are at least in part regulated through a previously unrecognized paracrine mechanism involving endothelial CEPT1 and hepatic MTTP signaling. Since selective knockdown of CEPT1 is essential for EC function ^14,18^, and selective hepatic knockdown of MTTP in mice is known to block the release of lipoproteins and leads to hepatic steatosis ^19,20^, we explored whether endothelial CEPT1 and hepatic MTTP had a molecular interaction that can impact hyperlipidemia and arterial atherosclerosis. We observed that human steatotic livers had suppressed MTTP activity, while there appeared to be decreased CEPT1 co-localization specifically in endothelial cells, but not in the liver overall. Selective, conditional knockdown of endothelial *Cept1 in vivo*, in the context of advanced hepatic steatosis, led to both a significant reduction in MTTP activity and a marked decrease in aortic atherosclerosis. The athero-protective impact of *Cept1* knockdown appeared to occur without evidence of fat malabsorption or weight loss, highlighting a unique hepatic regulatory axis involving the endothelium that is distinct from systemic lipid metabolism.

CEPT1 is essential for EC function and activity as we previously have shown ^13,14^. Our current study provides additional evidence that CEPT1 impacts EC signaling, particularly through paracrine signaling with hepatocytes. Our studies show that endothelial CEPT1 has functions that are beyond their traditional and established roles of phospholipogenesis and cell membrane homeaostasis (12,13). Specifically, we observed that *Cept1^fl/fl^Cre^+^Apoe^+/+^* mice with no hepatic steatosis had a significant reduction in hepatic MTTP activity, whereas *Cept1^fl/fl^Cre^+^Apoe^-/-^* mice with severe hepatic steatosis had no change in MTTP activity or protein content. These findings show that CEPT1 plays an important role in MTTP regulation even in settings where MTTP would be expected to be activated.

There are a few explanations for why MTTP activity is possibly reduced in *Cept1^fl/fl^Cre^+^Apoe^-/-^* mice. This may have resulted from heightened lipid biosynthesis, which could lead to antagonism of MTTP function and therefore obscure subtle functional alterations through metabolic compensation ^21–23^. While this interaction appears to be paracrine in nature, we observed direct effects on MTTP activity in hepatocytes when co-cultured with CEPT1- deficient ECs without physical contact. Another possible explanation is that reduced MTTP activity may stem from altered lipid trafficking and membrane composition in hepatocytes exposed to CEPT1-deficient endothelial signals. CEPT1 is critical for phosphatidylcholine synthesis, and its deficiency could perturb extracellular vesicle cargo or secreted lipid mediators that normally stabilize MTTP function (12,13). Such changes might impair the assembly of lipoproteins within hepatocytes, leading to apparent suppression of MTTP activity independent of transcriptional regulation. This mechanism highlights a potentially broader role of endothelial lipid metabolism in shaping hepatic lipoprotein handling.

Our study findings suggests that ECs in the liver may play an important role in regulating hepatic lipid transport ^5,24^. This adds to our understanding about endothelial-hepatocyte communication and its impact on lipid metabolism ^25^. *In vitro* studies utilizing a co-culture system of HUVECs and HepG2 cells strongly support the role of paracrine signaling as the mechanism supporting the interaction between the endothelium and hepatocyte function. This concept is supported by previous work that demonstrated that endothelial manipulation of lipid- handling proteins such as LRP1 (LDL Receptor-related Protein 1) can impact systemic lipid profiles ^26^, thus establishing a precedent to the broader role of ECs in regulating hepatic lipid metabolism via paracrine mechanisms. Our findings are also supported by literature demonstrating that liver sinusoidal ECs act as crucial paracrine regulators of hepatic physiology (20). When *Cept1* was silenced in HUVECs we observed reduced MTTP activity in co-cultured HepG2 cells. The physical separation of the ECs and hepatocytes in our co-culture system highlights the importance of secreted factors in mediating the observed effects. The associated reduction in *PPARα* expression in HepG2 cells, suggests that signaling mechanisms are blunting the transcription of factors that are downstream of PPARα. The endothelial-hepatocyte crosstalk highlights a lipid homeostasis signaling mechanism that appears to be distinct from what happens in ECs and hepatocytes individually ^29,30^.

MTTP plays a crucial role in regulatory pathways involved in lipoprotein assembly and secretion ^8,31,32^. The lipidation process facilitated by MTTP prevents ApoB degradation and enables the formation of mature VLDL particles in the liver, which are then transported via vesicular carriers from the ER to the Golgi apparatus for secretion into the circulation ^8^. Our study shows that MTTP appears to not only correlate with human disease severity but also seems to be regulated by CEPT1 and PPARα, establishing a previously unrecognized regulatory pathway ^33^. Unlike complete *Mttp* knockout, our endothelial *Cept1* knockdown in *Cept1^fl/fl^Cre^+^Apoe^+/+^* mice does not lead to a complete MTTP protein elimination; and only a slight decrease in lipid content via ORO staining when compared to *Cept1^fl/fl^Cre^-^Apoe^+/+^*. This observation suggests that endothelial CEPT1 is likely regulating MTTP-mediated VLDL assembly and lipid handling rather than directly affecting MTTP protein stability in hepatocytes.

The PPARα agonist fenofibrate is known to impact liver lipid metabolism and reduce serum triglyceride levels ^34–36^. Our study shows that fenofibrate can rescue MTTP activity, suggesting a clinically significant mechanism by which fenofibrate may influence circulating lipoprotein concentrations. ^37^. The fenofibrate mediated rescue of MTTP activity provides a plausible mechanism for the athero-protective phenotype observed in both preclinical rodent studies as well as in humans ^38^. Clinical trial data further support the impact of fenofibrate on hepatic steatosis. For example, a pilot trial in patients with biopsy-confirmed NAFLD (Non- Alcoholic Fatty Liver Disease) found that fenofibrate treatment improved biochemical and histological markers of liver disease ^39^. Other clinical studies have shown that fenofibrate monotherapy can reduce hepatic steatosis in patients with metabolic syndrome and ultrasonographic evidence of NAFLD ^40^.

Furthermore, our observations of paracrine effects between the endothelium and hepatic lipid metabolism are reinforced by a study on PPARα in setting of hepatocellular carcinoma ^41^. Prior findings show that endothelial PPARα activation can limit mitochondrial oxidative stress and inhibit inflammatory signaling in liver tissue, supporting our hypothesis of endothelial- hepatocyte crosstalk in lipid regulation ^42^. While prior work linked endothelial PPARγ to FA uptake regulation, our data implicates CEPT1 in a distinct PPARα-MTTP molecular axis, suggesting it as a potential novel target for regulation of lipid disorders and hyperlipidemia ^43,44^.

The therapeutic potential of targeting the endothelium is underscored by our observations showing conditional *Cept1* knockdown significantly reduced serum lipid levels and aortic plaque area. Notably, endothelial CEPT1 knockout achieved a ∼70% reduction in aortic lesion area, surpassing the 24–60% reductions reported for targeting other types of proteins such as CCR2, CX3CR1, IFN-γ receptor, ICAM-1, P-selectin, or E-selectin in *Apoe^-/-^* mice ^29(p),45–48^. Importantly, the athero-protective phenotype observed in mice with *Cept1* knockdown occurs without evidence of gastrointestinal fat malabsorption or weight loss. This further distinguishes the impact of CEPT1 on systemic lipid metabolism since it does not appear to have a detectable deleterious systemic side effect ^49^. Therefore, our findings make endothelial CEPT1 an attractive target that can impact atheroprogression without dramatically disrupting normal metabolic functions. For example, we previously demonstrated that endothelial-specific knockdown of *Cept1* in mice resulted in reduced perfusion and angiogenesis in ischemic limbs, but fenofibrate (a PPARα agonist) can rescue these effects, suggesting that modulating CEPT1- related signaling pathways may be both effective and reversible ^50^.

Our findings also reveal that hepatic steatosis influences CEPT1 expression in the endothelium, suggesting a signaling mechanism linking liver disease to vascular dysfunction and disease progression ^51,52^. Emerging evidence indicates that liver-derived signals such as extracellular vesicles and altered endothelial gene expression mediate this relationship, linking liver fat accumulation to endothelial dysfunction and accelerated atherosclerosis ^53–55^. Notably, *Cept1* knockout mice exhibited reduced serum cholesterol and triglycerides, as well as less severe aortic atherosclerosis ^14^. This is supported by observations that *Cept1* knockdown in the liver leads to alterations in lipid uptake, *de novo* lipogenesis, β-oxidation, or lipid secretion. Earlier studies show that CEPT1 is elevated in diseased human arteries and that its loss reduces EC proliferation, migration, and recovery after injury ^50^. This effect relies on PPARα signaling and is linked to liver and vascular health ^56–59^. Future studies should continue to focus investigation on how CEPT1 and MTTP signaling mechanisms can be targeted to reduce the risk of hepatic steatosis and thereby reduce cardiovascular complications and improve overall patient outcomes ^60^.

In conclusion, our findings advance the understanding of EC metabolism in atherosclerosis by identifying CEPT1 as a key regulator of atheroprogression and hepatic steatosis ^61^. We demonstrate that ECs can influence hepatocyte lipid metabolism and atherosclerosis progression ^62^. Our findings offer new avenues for targeted interventions in atherosclerosis and related metabolic disorders, potentially leading to more effective strategies for cardiovascular disease prevention and treatment ^63^. The identification of endothelial CEPT1 as a potential therapeutic target may lead to the development of novel pharmacological approaches that selectively reduce atherosclerosis without disrupting normal metabolic functions^64^.

## Acknowledgments

This study was supported by the vascular biobank at Washington University School of Medicine in St. Louis.

## Non-standard Abbreviations and Acronyms

ApoB: Apolipoprotein B
BSA: Bovine Serum Albumin
CCR2: C-C Chemokine Receptor Type 2
CEPT1: Choline Ethanolamine Phosphotransferase 1
DAPI: 4’,6-Diamidino-2-Phenylindole
EC: Endothelial Cell
FEN: Fenofibrate
FFA: Free Fatty Acid
GAPDH: Glyceraldehyde 3-Phosphate Dehydrogenase
HFD: High-Fat Diet
HRPO: Human Research Protection Office
HUVEC: Human Umbilical Vein Endothelial Cell
IACUC: Institutional Animal Care and Use Committee
IFN-γ: Interferon Gamma
LRP1: LDL Receptor-related Protein 1
MTTP: Microsomal Triglyceride Transfer Protein
NAFLD: Non-Alcoholic Fatty Liver Disease
ORO: Oil Red O
PC: Phosphatidylcholine
PE: Phosphatidylethanolamine
PPAR: Peroxisome Proliferator-Activated Receptor
PVDF: Polyvinylidene Difluoride
RT-PCR: Real-Time Polymerase Chain Reaction
TBS-T: Tris-Buffered Saline with Tween 20
VLDL: Very-Low-Density Lipoprotein

## Supplementary Figure Legends

**Supplementary Figure 1. Expression of *MTTP* and *CEPT1* mRNA in human liver tissue.** RT-PCR analysis of MTTP and CEPT1 mRNA levels in liver tissue samples from individuals with non-steatosis (n =6) and steatosis (n =6). Data are presented as mean ± SEM (or SD, as appropriate) and normalized to a housekeeping gene (GAPDH).

**Supplementary Figure 2. *In Vivo* fecal lipid measurements.** (A–C) Fecal cholesterol (A), triglycerides (B), and free fatty acids (FFA; C) measured in *Cept1^fl/fl^Cre^-^Apoe^+/+^*and *Cept1^fl/fl^Cre^+^Apoe^+/+^* mice at 6, 10, 14, and 18 weeks on high-fat diet (HFD). (D–F) Fecal cholesterol (D), triglycerides (E), and free fatty acids (FFA; F) measured in *Cept1^fl/fl^Cre^-^Apoe^-/-^*and *Cept1^fl/fl^Cre^+^Apoe^-/-^* mice at at the indicated time points. Data are presented as μg/mg dry fecal weight.

**Supplementary Figure 3. Hepatic lipid levels in endothelial-specific *Cept1* knockout ice.** Quantification of hepatic (A) triglycerides, (B) total cholesterol, and (C) free fatty acids normalized to protein content in liver homogenates from *Cept1^fl/fl^Cre^+^Apoe^-/-^*and *Cept1^fl/fl^Cre^+^Apoe^+/+^*versus corresponding controls (n=3).

**Supplementary Figure 4. Endothelial-specific *Cept1* knockdown leads to reduced atherosclerotic plaque in the aortic arch.** (A) Representative examples of atherosclerotic plaque (yellow arrows) in the aortic arches of *Cept1^fl/fl^Cre^+^Apoe^-/-^* and *Cept1^fl/fl^Cre^+^Apoe^-/-^* mice (2 mice per genotype). Scale bar=1cm.

## References

1. Ghazwani M, Mahmood SE, Gosadi IM, Bahri AA, Ghazwani SH, Khmees RA. Prevalence of Dyslipidemia and Its Determinants Among the Adult Population of the Jazan Region. IJGM. 2023;16:4215–4226. doi:10.2147/IJGM.S429462

2. Mohamed-Yassin MS, Baharudin N, Abdul-Razak S, Ramli AS, Lai NM. Global prevalence of dyslipidaemia in adult populations: a systematic review protocol. BMJ Open. 2021;11(12):e049662. doi:10.1136/bmjopen-2021-049662

3. Younossi ZM, Koenig AB, Abdelatif D, Fazel Y, Henry L, Wymer M. Global epidemiology of nonalcoholic fatty liver disease-Meta-analytic assessment of prevalence, incidence, and outcomes. Hepatology. 2016;64(1):73–84. doi:10.1002/hep.28431

4. Pi X, Xie L, Patterson C. Emerging Roles of Vascular Endothelium in Metabolic Homeostasis. Circulation Research. 2018;123(4):477–494. doi:10.1161/CIRCRESAHA.118.313237

5. Kim B, Arany Z. Endothelial Lipid Metabolism. Cold Spring Harb Perspect Med. 2022;12(6):a041162. doi:10.1101/cshperspect.a041162

6. Goldberg IJ, Bornfeldt KE. Lipids and the Endothelium: Bidirectional Interactions. Curr Atheroscler Rep. 2013;15(11):10.1007/s11883-013-0365-1. doi:10.1007/s11883-013-0365-1

7. Iqbal J, Jahangir Z, Al-Qarni AA. Microsomal Triglyceride Transfer Protein: From Lipid Metabolism to Metabolic Diseases. In: Jiang XC, ed. Lipid Transfer in Lipoprotein Metabolism and Cardiovascular Disease. Springer; 2020:37–52. doi:10.1007/978-981-15-6082-8_4

8. Hussain MM, Shi J, Dreizen P. Microsomal triglyceride transfer protein and its role in apoB-lipoprotein assembly. J Lipid Res. 2003;44(1):22–32. doi:10.1194/jlr.r200014-jlr200

9. Raabe M, Véniant MM, Sullivan MA, et al. Analysis of the role of microsomal triglyceride transfer protein in the liver of tissue-specific knockout mice. J Clin Invest. 1999;103(9):1287–1298.

10. Skogsberg J, Lundström J, Kovacs A, et al. Transcriptional Profiling Uncovers a Network of Cholesterol-Responsive Atherosclerosis Target Genes. PLOS Genetics. 2008;4(3):e1000036. doi:10.1371/journal.pgen.1000036

11. Hewing B, Parathath S, Mai CK, Fiel MI, Guo L, Fisher EA. Rapid regression of atherosclerosis with MTP inhibitor treatment. Atherosclerosis. 2013;227(1):125–129. doi:10.1016/j.atherosclerosis.2012.12.026

12. Wang Z, Yang M, Yang Y, He Y, Qian H. Structural basis for catalysis of human choline/ethanolamine phosphotransferase 1. Nat Commun. 2023;14(1):2529. doi:10.1038/s41467-023-38290-2

13. Dorighello G, McPhee M, Halliday K, Dellaire G, Ridgway ND. Differential contributions of phosphotransferases CEPT1 and CHPT1 to phosphatidylcholine homeostasis and lipid droplet biogenesis. J Biol Chem. 2023;299(4):104578. doi:10.1016/j.jbc.2023.104578

14. Zayed MA, Jin X, Yang C, et al. CEPT1-Mediated Phospholipogenesis Regulates Endothelial Cell Function and Ischemia-Induced Angiogenesis Through PPARα. Diabetes. 2021;70(2):549–561. doi:10.2337/db20-0635

15. Edvardsson U, Ljungberg A, Lindén D, et al. PPARα activation increases triglyceride mass and adipose differentiation-related protein in hepatocytes. Journal of Lipid Research. 2006;47(2):329–340. doi:10.1194/jlr.M500203-JLR200

16. Sun W, Cui B, Hong F, Xu Y. Establishment of ApoE-knockout mouse model of preeclampsia and relevant mechanisms. Exp Ther Med. 2016;12(4):2634–2638. doi:10.3892/etm.2016.3678

17. Meade R, Ibrahim D, Engel C, et al. Targeting fatty acid synthase reduces aortic atherosclerosis and inflammation. Communications Biology. 2025;8(1):262. doi:10.1038/s42003-025-07656-1

18. Hafezi S, Khan TJ, Belaygorod L, Arif B, Semenkovich CF, Zayed MA. Endothelial CEPT1 Impacts Aortic Atherosclerosis and Hepatic Lipoprotein Metabolism. JVS-Vascular Science. 2024;5. doi:10.1016/j.jvssci.2024.100234

19. Newberry EP, Xie Y, Kennedy SM, et al. Prevention of hepatic fibrosis with liver microsomal triglyceride transfer protein deletion in liver fatty acid binding protein null mice. Hepatology. 2017;65(3):836–852. doi:10.1002/hep.28941

20. Hooper AJ, Burnett JR, Watts GF. Contemporary Aspects of the Biology and Therapeutic Regulation of the Microsomal Triglyceride Transfer Protein. Circulation Research. 2015;116(1):193–205. doi:10.1161/CIRCRESAHA.116.304637

21. Wagner T, Bartelt A, Schlein C, Heeren J. Genetic Dissection of Tissue-Specific Apolipoprotein E Function for Hypercholesterolemia and Diet-Induced Obesity. PLoS One. 2015;10(12):e0145102. doi:10.1371/journal.pone.0145102

22. Schierwagen R, Maybüchen L, Zimmer S, et al. Seven weeks of Western diet in apolipoprotein-E-deficient mice induce metabolic syndrome and non-alcoholic steatohepatitis with liver fibrosis. Sci Rep. 2015;5(1):12931. doi:10.1038/srep12931

23. Zhao J, Liu X, Yue J, Zhang S, Li L, Wei H. PF-05231023 reduces lipid deposition in apolipoprotein E-deficient mice by inhibiting the expression of lipid synthesis genes. Front Vet Sci. 2024;11. doi:10.3389/fvets.2024.1429639

24. Du W, Wang L. The Crosstalk Between Liver Sinusoidal Endothelial Cells and Hepatic Microenvironment in NASH Related Liver Fibrosis. Front Immunol. 2022;13:936196. doi:10.3389/fimmu.2022.936196

25. Sun X, Harris EN. New aspects of hepatic endothelial cells in physiology and nonalcoholic fatty liver disease. Am J Physiol Cell Physiol. 2020;318(6):C1200–C1213. doi:10.1152/ajpcell.00062.2020

26. Mao H, Lockyer P, Li L, et al. Endothelial LRP1 regulates metabolic responses by acting as a co-activator of PPARγ. Nat Commun. 2017;8(1):14960. doi:10.1038/ncomms14960

27. Rowe IA, Galsinh SK, Wilson GK, et al. Paracrine Signals From Liver Sinusoidal Endothelium Regulate Hepatitis C Virus Replication. Hepatology. 2014;59(2):375–384. doi:10.1002/hep.26571

28. DeLeve LD, Wang X, Hu L, McCuskey MK, McCuskey RS. Rat liver sinusoidal endothelial cell phenotype is maintained by paracrine and autocrine regulation. Am J Physiol Gastrointest Liver Physiol. 2004;287(4):G757–763. doi:10.1152/ajpgi.00017.2004

29. Bougarne N, Weyers B, Desmet SJ, et al. Molecular Actions of PPARα in Lipid Metabolism and Inflammation. Endocrine Reviews. 2018;39(5):760–802. doi:10.1210/er.2018-00064

30. Bort A, Sánchez BG, Mateos-Gómez PA, Díaz-Laviada I, Rodríguez-Henche N. Capsaicin Targets Lipogenesis in HepG2 Cells Through AMPK Activation, AKT Inhibition and PPARs Regulation. Int J Mol Sci. 2019;20(7):1660. doi:10.3390/ijms20071660

31. Hussain MM, Rava P, Walsh M, Rana M, Iqbal J. Multiple functions of microsomal triglyceride transfer protein. Nutrition & Metabolism. 2012;9(1):14. doi:10.1186/1743-7075-9-14

32. Iqbal J, Jahangir Z, Al-Qarni AA. Microsomal Triglyceride Transfer Protein: From Lipid Metabolism to Metabolic Diseases. Adv Exp Med Biol. 2020;1276:37–52. doi:10.1007/978-981-15-6082-8_4

33. Améen C, Edvardsson U, Ljungberg A, et al. Activation of peroxisome proliferator-activated receptor alpha increases the expression and activity of microsomal triglyceride transfer protein in the liver. J Biol Chem. 2005;280(2):1224–1229. doi:10.1074/jbc.M412107200

34. Lakhia R, Yheskel M, Flaten A, Quittner-Strom EB, Holland WL, Patel V. PPARα agonist fenofibrate enhances fatty acid β-oxidation and attenuates polycystic kidney and liver disease in mice. Am J Physiol Renal Physiol. 2018;314(1):F122–F131. doi:10.1152/ajprenal.00352.2017

35. Nikam A, Patankar JV, Somlapura M, et al. The PPARα Agonist Fenofibrate Prevents Formation of Protein Aggregates (Mallory-Denk bodies) in a Murine Model of Steatohepatitis-like Hepatotoxicity. Sci Rep. 2018;8(1):12964. doi:10.1038/s41598-018-31389-3

36. Yoo J, Jeong IK, Ahn KJ, Chung HY, Hwang YC. Fenofibrate, a PPARα agonist, reduces hepatic fat accumulation through the upregulation of TFEB-mediated lipophagy. Metabolism. 2021;120:154798. doi:10.1016/j.metabol.2021.154798

37. Keech A, Simes RJ, Barter P, et al. Effects of long-term fenofibrate therapy on cardiovascular events in 9795 people with type 2 diabetes mellitus (the FIELD study): randomised controlled trial. Lancet. 2005;366(9500):1849-1861. doi:10.1016/S0140-6736(05)67667-2

38. Rubins HB, Robins SJ, Collins D, et al. Gemfibrozil for the Secondary Prevention of Coronary Heart Disease in Men with Low Levels of High-Density Lipoprotein Cholesterol. New England Journal of Medicine. 1999;341(6):410–418. doi:10.1056/NEJM199908053410604

39. Fernández-Miranda C, Pérez-Carreras M, Colina F, López-Alonso G, Vargas C, Solís- Herruzo JA. A pilot trial of fenofibrate for the treatment of non-alcoholic fatty liver disease. Digestive and Liver Disease. 2008;40(3):200–205. doi:10.1016/j.dld.2007.10.002

40. Kostapanos MS, Kei A, Elisaf MS. Current role of fenofibrate in the prevention and management of non-alcoholic fatty liver disease. World J Hepatol. 2013;5(9):470–478. doi:10.4254/wjh.v5.i9.470

41. Pawlak M, Lefebvre P, Staels B. Molecular mechanism of PPARα action and its impact on lipid metabolism, inflammation and fibrosis in non-alcoholic fatty liver disease. Journal of Hepatology. 2015;62(3):720–733. doi:10.1016/j.jhep.2014.10.039

42. Ip E, Farrell GC, Robertson G, Hall P, Kirsch R, Leclercq I. Central role of PPARalpha- dependent hepatic lipid turnover in dietary steatohepatitis in mice. Hepatology. 2003;38(1):123–132. doi:10.1053/jhep.2003.50307

43. Leone TC, Weinheimer CJ, Kelly DP. A critical role for the peroxisome proliferator-activated receptor alpha (PPARalpha) in the cellular fasting response: the PPARalpha-null mouse as a model of fatty acid oxidation disorders. Proc Natl Acad Sci U S A. 1999;96(13):7473–7478. doi:10.1073/pnas.96.13.7473

44. Rakhshandehroo M, Knoch B, Müller M, Kersten S. Peroxisome proliferator-activated receptor alpha target genes. PPAR Res. 2010;2010:612089. doi:10.1155/2010/612089

45. Boring L, Gosling J, Cleary M, Charo IF. Decreased lesion formation in CCR2−/− mice reveals a role for chemokines in the initiation of atherosclerosis. Nature. 1998;394(6696):894-897. doi:10.1038/29788

46. Combadière C, Potteaux S, Gao JL, et al. Decreased atherosclerotic lesion formation in CX3CR1/apolipoprotein E double knockout mice. Circulation. 2003;107(7):1009–1016. doi:10.1161/01.cir.0000057548.68243.42

47. Gupta S, Pablo AM, Jiang X c, Wang N, Tall AR, Schindler C. IFN-gamma potentiates atherosclerosis in ApoE knock-out mice. J Clin Invest. 1997;99(11):2752–2761.

48. Geng JG, Chen M, Chou KC. P-selectin Cell Adhesion Molecule in Inflammation, Thrombosis, Cancer Growth and Metastasis. Current Medicinal Chemistry. 2004;11(16):2153–2160. doi:10.2174/0929867043364720

49. Wetterau JR, Gregg RE, Harrity TW, et al. An MTP Inhibitor That Normalizes Atherogenic Lipoprotein Levels in WHHL Rabbits. Science. 1998;282(5389):751-754. doi:10.1126/science.282.5389.751

50. Zayed MA, Jin X, Yang C, et al. CEPT1-Mediated Phospholipogenesis Regulates Endothelial Cell Function and Ischemia-Induced Angiogenesis Through PPARα. Diabetes. 2021;70(2):549–561. doi:10.2337/db20-0635

51. Francque S, Verrijken A, Mertens I, et al. Visceral adiposity and insulin resistance are independent predictors of the presence of non-cirrhotic NAFLD-related portal hypertension. Int J Obes (Lond). 2011;35(2):270–278. doi:10.1038/ijo.2010.134

52. Francque S, Wamutu S, Chatterjee S, et al. Non-alcoholic steatohepatitis induces non-fibrosis-related portal hypertension associated with splanchnic vasodilation and signs of a hyperdynamic circulation in vitro and in vivo in a rat model. Liver Int. 2010;30(3):365–375. doi:10.1111/j.1478-3231.2009.02136.x

53. Chen X, Chen S, Pang J, et al. Hepatic steatosis aggravates atherosclerosis via small extracellular vesicle-mediated inhibition of cellular cholesterol efflux. J Hepatol. 2023;79(6):1491–1501. doi:10.1016/j.jhep.2023.08.023

54. Nasiri-Ansari N, Androutsakos T, Flessa CM, et al. Endothelial Cell Dysfunction and Nonalcoholic Fatty Liver Disease (NAFLD): A Concise Review. Cells. 2022;11(16):2511. doi:10.3390/cells11162511

55. 55. Gato S, García-Fernández V, Gil-Gómez A, et al. Navigating the Link Between Non-alcoholic Fatty Liver Disease/Non-alcoholic Steatohepatitis and Cardiometabolic Syndrome. Published online December 27, 2023. Accessed June 16, 2025. https://www.ecrjournal.com/articles/navigating-link-between-non-alcoholic-fatty-liver-diseasenon-alcoholic-steatohepatitis-and?language_content_entity=en

56. Donnelly KL, Smith CI, Schwarzenberg SJ, Jessurun J, Boldt MD, Parks EJ. Sources of fatty acids stored in liver and secreted via lipoproteins in patients with nonalcoholic fatty liver disease. J Clin Invest. 2005;115(5):1343–1351. doi:10.1172/JCI23621

57. Postic C, Girard J. Contribution of de novo fatty acid synthesis to hepatic steatosis and insulin resistance: lessons from genetically engineered mice. J Clin Invest. 2008;118(3):829–838. doi:10.1172/JCI34275

58. Fabbrini E, Mohammed BS, Magkos F, Korenblat KM, Patterson BW, Klein S. Alterations in adipose tissue and hepatic lipid kinetics in obese men and women with nonalcoholic fatty liver disease. Gastroenterology. 2008;134(2):424–431. doi:10.1053/j.gastro.2007.11.038

59. Lambert JE, Ramos-Roman MA, Browning JD, Parks EJ. Increased de novo lipogenesis is a distinct characteristic of individuals with nonalcoholic fatty liver disease. Gastroenterology. 2014;146(3):726–735. doi:10.1053/j.gastro.2013.11.049

60. Anstee QM, Targher G, Day CP. Progression of NAFLD to diabetes mellitus, cardiovascular disease or cirrhosis. Nat Rev Gastroenterol Hepatol. 2013;10(6):330–344. doi:10.1038/nrgastro.2013.41

61. Ding BS, Nolan DJ, Butler JM, et al. Inductive angiocrine signals from sinusoidal endothelium are required for liver regeneration. Nature. 2010;468(7321):310-315. doi:10.1038/nature09493

62. Hu J, Srivastava K, Wieland M, et al. Endothelial cell-derived angiopoietin-2 controls liver regeneration as a spatiotemporal rheostat. Science. 2014;343(6169):416-419. doi:10.1126/science.1244880

63. Poisson J, Lemoinne S, Boulanger C, et al. Liver sinusoidal endothelial cells: Physiology and role in liver diseases. J Hepatol. 2017;66(1):212–227. doi:10.1016/j.jhep.2016.07.009

64. DeLeve LD. Liver sinusoidal endothelial cells in hepatic fibrosis. Hepatology. 2015;61(5):1740–1746. doi:10.1002/hep.27376

